# Nucleated Fish Erythrocyte Extracellular Traps (FEETs) release is an evolutionary conserved immune defence process

**DOI:** 10.1101/2021.08.10.455826

**Authors:** Giulia Rinaldi, Neila Álvarez de Haro, Andrew P. Desbois, Calum T. Robb, Adriano G. Rossi

## Abstract

Fish erythrocytes remain nucleated for their life-span, unlike mammalian erythrocytes which undergo enucleation. Asides transportation of oxygen, fish erythrocytes are capable of several immune defence processes. Nucleated fish erythrocytes represent prime candidates for carrying out ETotic responses. ETosis is an evolutionary conserved innate immune defence process found in both vertebrates and invertebrates, which involves the extrusion of DNA studded with antimicrobial proteins into the extracellular space serving to trap and kill microorganisms. In this report, we demonstrate that fish erythrocytes isolated from *Danio rerio* (zebrafish) produce ETotic-like responses when exposed to chemical and physiological stimuli. Furthermore, we found *Salmo salar* (Atlantic salmon) erythrocytes produce similar ETotic responses. We have termed these ET-like formations Fish Erythrocyte Extracellular Traps (FEETs). Interestingly, we discovered that mammalian inducers of NETosis, such as the protein kinase C (PKC) activator phorbol 12-myristate 13-acetate and the calcium ionophore ionomycin, induced FEETs. Moreover, we found that FEETs are dependent upon activation of PKC and generation of mitochondrial reactive oxygen species. Thus, this brief report represents the first demonstration that fish erythrocytes can exhibit ETotic-like responses, unveiling a previously unknown function of nucleated erythrocytes, which sheds new light on the innate immune arsenal of erythrocytes.

## Introduction

Mammalian erythrocytes carry oxygen to tissues and, during erythropoiesis, prior to entering the circulation, undergo ‘enucleation’ where the nucleus is extruded^1^. In contrast, certain organisms, including fish, possess erythrocytes that remain nucleated for their life-span and contain cytoplasmic organelles^2^. Asides from oxygen carriage, erythrocytes are capable of several immune functions such as antigen presentation, phagocytosis, generation of reactive oxygen species (ROS) and mediator production than can inhibit macrophage migration^3^. Furthermore, these cells are transcriptionally active, allowing upregulation of immune-related genes in response to invading pathogens^4,5^. Such evidence supports our hypothesis that erythrocytes can function somewhat similarly to leukocytes and play an important role in immune defence.

Extracellular trap (ET) formation is an immune defence response first described in human neutrophils (neutrophil extracellular traps (NETs))^6^ whereby nuclear material is released during a process termed NETosis. This nuclear material is studded with granular products which, together with DNA, histones and antimicrobial proteins, serve to entrap and neutralise invading organisms such as bacteria and fungi. This phenomenon occurs in other mammalian immune cells including monocytes^7^, macrophages^8^, eosinophils^9^ and mast cells^10^, leading to the collective term of ETosis^11^. ETosis is also exhibited by immune cells of non-mammalian vertebrates and invertebrates^12,13^ and is an evolutionary conserved immune defence mechanism^13^. However, mammalian ETosis is viewed as a double-edged sword. ETosis-mediated entrapment and neutralisation of microorganisms is essential, conferring highly localised antimicrobial activity, initiation of coagulation cascades, as well as anti-inflammatory activities. However, aberrant ETosis is implicated in a wide range of ETosis-associated diseases including correlation of ETs and severe SARS-CoV-2 conditions^14,15,16^. In fish, ETosis occurs in neutrophils^17,18^ and macrophages^19^ of several species, including zebrafish (*Danio rerio*) ^20^ and rainbow trout (*Oncorhynchus mykiss*)^18^. Here, we hypothesised that nucleated fish erythrocytes exhibit evolutionally conserved ETotic-like responses.

We provide the first evidence of ETotic-like formation in nucleated fish erythrocytes in response to established mammalian inducers of NETs, including the protein kinase C activator phorbol 12-myristate 13-acetate (PMA), the calcium ionophore ionomycin and the fungus *Candida albicans*. *Such structures were* termed Fish Erythrocyte Extracellular Traps (FEETs). We observe that both zebrafish and Atlantic salmon (*Salmo salar*) erythrocytes exhibit FEETs distinctly similar in morphology to mammalian NETs ^6,21^. Using zebrafish as our main model system, we observed and quantified FEETs induced by PMA, calcium ionophore ionomycin and *C. albicans*. In addition, we report that PMA-induced FEET production is PKC dependent and ionomycin-induced FEET generation is mitochondrial ROS-dependent akin to mammalian NETs^21,22^. These novel findings shed new light on the potential beneficial roles that nucleated erythrocytes could play during non-mammalian vertebrate immunity.

## Study design

Blood was collected from zebrafish prior to euthanasia by retro-orbital (RO) collection^23^ and methods for Atlantic salmon can be found in Supplementary Material). All experiments were approved and carried out in a UK Home Office approved facility at the University of Edinburgh under a UK Home Office Project Licence in accordance with the European Directive 63/2010/EU. The salmon experiments were approved by the Ethics Committee of the Institute of Aquaculture at the University of Stirling. All the experiments were performed in accordance with animal euthanasia schedule 1 procedure (Animal Act 1986). Differential cell counts were performed by analysis of cytocentrifuge preparations to determine the purity of erythrocyte populations (detailed method in supplementary information). Viability and FEET production were quantified by culturing erythrocytes in 24 or 96 well flat-bottom plates (Costar) at 28.5°C (zebrafish) or 15°C (Atlantic salmon) ± established ET inducers (see below), and stained with SYTOX Green nucleic acid stain (1 *μ*M; 20 min)(Thermo Fisher Scientific). Membrane integrity was assessed from SYTOX Green positive (membrane integrity lost) and negative (membrane integrity retained) cells. ETs were quantified as percentage of cells forming ETs compared to the total number of cells in the fields of view (a minimum of 4 images from 4 different fields of view per technical replication at 20x objective lens were counted for each condition and experiment). FEETs were induced with established ETosis inducers, PMA (Sigma)^6,21^ and ionomycin (Sigma)^24^. To investigate if FEETs are produced via generation of cytosolic ROS, diphenylene iodonium (DPI; Sigma)^21^, and the mitochondrial ROS uncoupler, 2,4 dinitrophenol (DNP; Sigma) ^22^ were used. To further investigate if FEETs are generated via activation of PKC and activation of small conductance calcium-activated potassium (SK) channels, the established pan-PKC inhibitor Ro 31-8220 (Santa Cruz Biotechnology)^25^ and SK channel inhibitor apamin (Sigma)^22^ were used. Erythrocytes were co-cultured for various timepoints in L-15 Leibovitz’s media (28.5°C) with stimulators and/or inhibitors, before staining and imaging as above. FEETs were quantified by differential counts as described previous. To confirm the DNA composition of FEETs, DNase-I (Thermo Fisher Scientific) was added to digest extracellular DNA and images taken from the same field of view at regular time intervals. The Gram-positive bacterium *Staphylococcus aureus*, and the fungus *C. albicans*, were also used in attempt to stimulate FEETs. *S. aureus* was cultured in Luria-Bertani (LB) broth and heat-killed by thermal treatment (90°C; 1h). *C. albicans* was cultured in potato dextrose broth and heat-killed similarly. Each microbe was washed twice and resuspended in L-15 media at the desired concentration and co-cultured with zebrafish erythrocytes at various multiplicity of infections (MOIs).

## Results and discussion

First, we assessed zebrafish erythrocyte purity and established optimal culturing conditions. We found that >98% of circulating blood cells were erythrocytes (**Figure 1A i, ii**). To optimise experimental culturing times in zebrafish, we assessed erythrocyte membrane integrity by SYTOX Green staining over 48h (**Figure 1A iii**). There was no statistically significant difference when comparing membrane integrity retained at 1h or 3h *vs* 24h or 48h of culture (**Figure 1A iii, iv**). PMA- and ionomycin-induced human NETs exhibit classical NETotic diffuse and spread morphology (**Figure 1B iii**) and similar morphologies were exhibited by FEETs (**Figure 1B ii**). Different concentrations of established ETosis inducers were used in an attempt to stimulate FEETs at different time-points (**Figure 1B i**). We discovered that zebrafish erythrocytes released FEETs in response to PMA and ionomycin following 24h culture (**Figure 1B i,ii merge, white arrows**). Atlantic salmon showed similar significant increases in ETotic responses when stimulated by ionomycin (**Figure S3 B,C**). A significant increase in zebrafish FEET release was observed with PMA 10 *μ*M after 24 h of culture compared to 1h. The same effect was observed in response to 1 *μ*M and 10 *μ*M ionomycin at 1h vs 24h. Moreover, heat shock-induced zebrafish FEETs were observed by culturing erythrocytes at higher temperature (37 °C; 4 h)(**Figure 1B ii merge, white arrows**).

**Figure 1.**
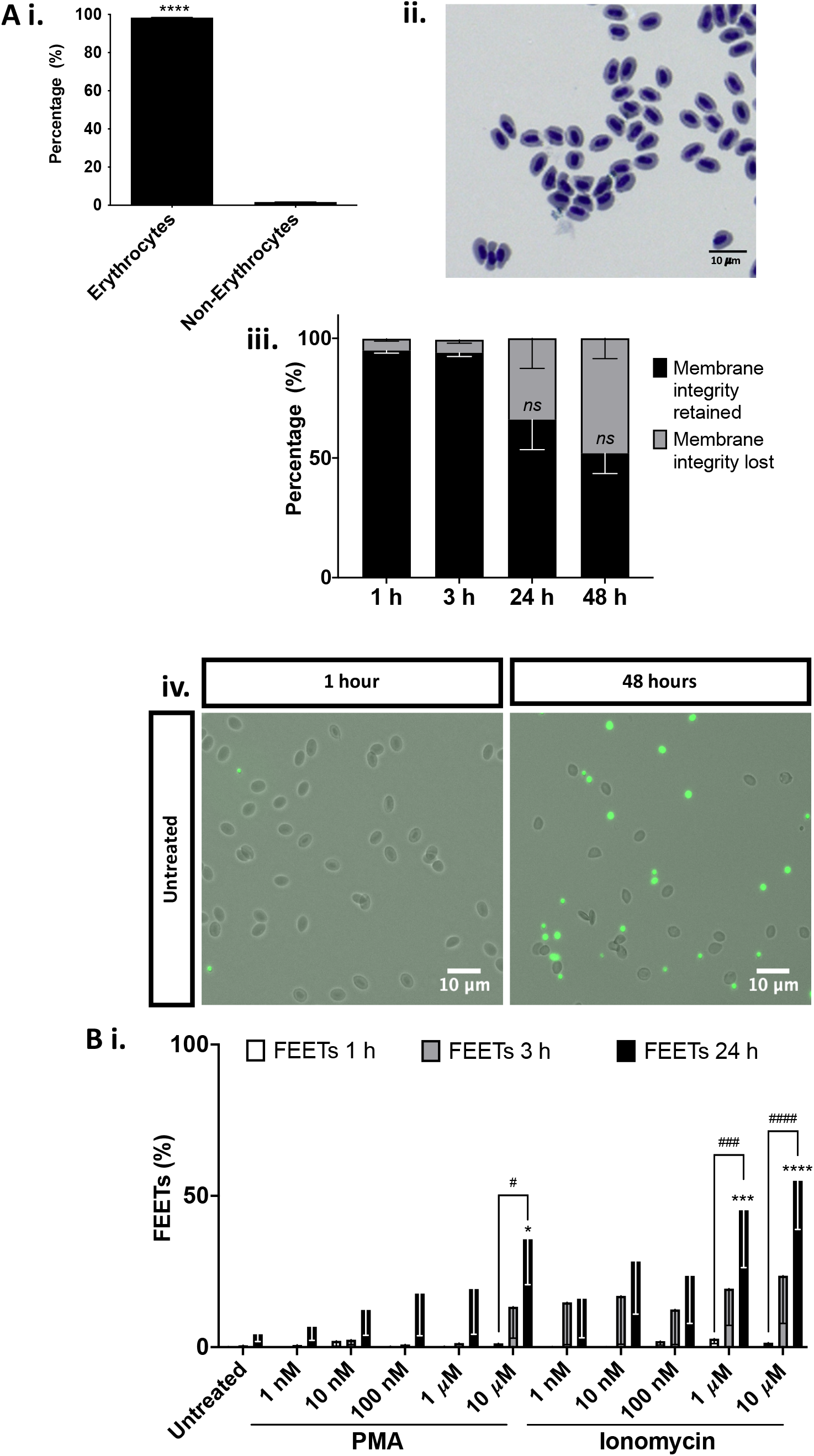
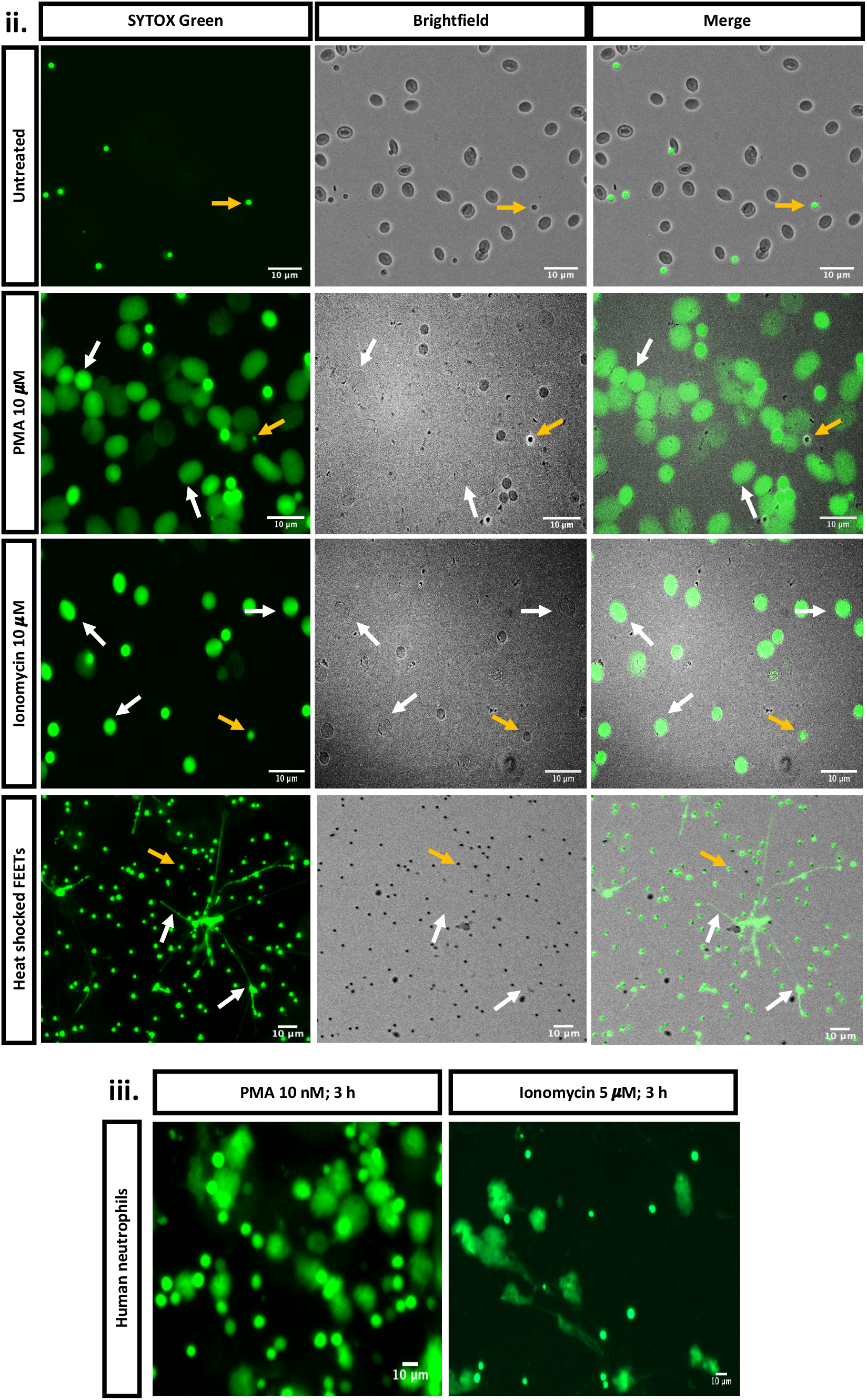
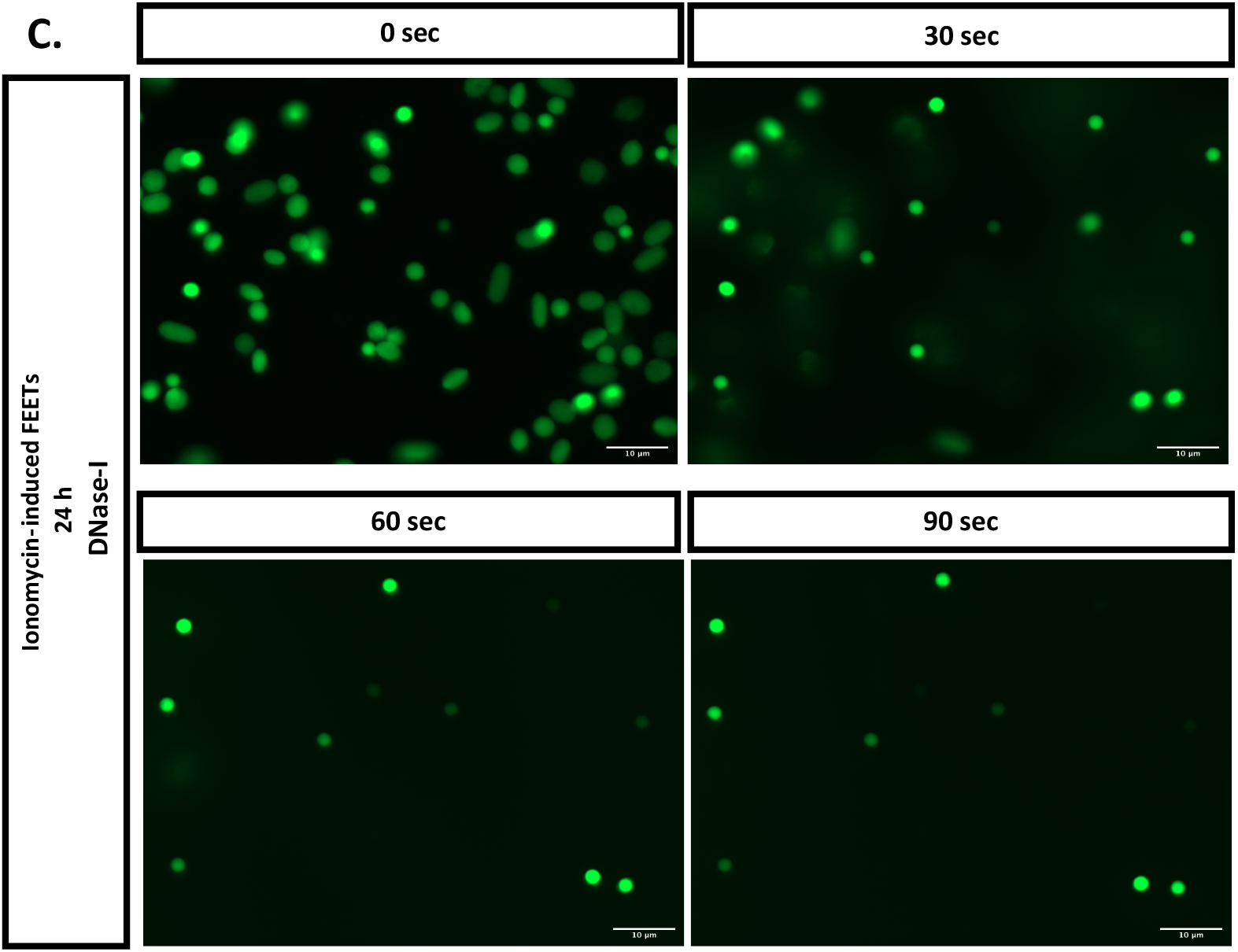
Zebrafish erythrocyte purity, membrane integrity and formation of FEETs in response to PMA and ionomycin. (**A i**): Zebrafish erythrocyte purity assessment after RO cavity blood collection quantified via differential counts of cytocentrifuge preparations. Values are expressed as mean percentage ± s.e.m., n=3. Significant difference between erythrocytes and non-erythrocytes (**** *P*<0.0001) calculated by a Paired t-test. (**A ii**): Representative cytocentrifuge preparation image of zebrafish whole blood. (**A iii**): Untreated erythrocyte membrane integrity retention after 1, 3, 6, 24 and 48 h culture in L-15 media and quantified via differential counts by SYTOX Green staining (1 *μ*M; 30 min). Values are expressed as mean percentage ± s.e.m., n=6. Comparison between 1 h or 3 h membrane integrity retained vs 24 or 48 h did not reach significance (*ns P*>0.05) calculated by 2-way ANOVA analysis with Tukey’s multiple comparison *post hoc* test. (**A iv**): Representative brightfield and GFP merge images of untreated erythrocytes after 1 h (left) and 48 h (right) culture (28°C; L-15 media). Images taken under 20x objective lens, scale bar 10 *μ*m. (**B i**): ET formations in erythrocytes following PMA or Ionomycin treatment (1, 3 or 24 h) at different concentrations (1 nM, 10 nM, 100 nM, 1 *μ*M or 10 *μ*M) or left untreated (control). Values expressed as mean percentage ± s.e.m., n=6. Significant differences between PMA or ionomycin treatment and untreated (control) (* = Untreated 24 h *vs* PMA 10 *μ*M 24 h, *P*<0.05; ***= Untreated 24 h *vs* ionomycin 1 *μ*M 24 h, *P*<0.001; **** = Untreated 24 h *vs* ionomycin 10 *μ*M 24 h, *P*<0.0001). Significant differences across time points of PMA and ionomycin treatment (# = PMA 10 *μ*M 1 h *vs* PMA 10 *μ*M 24 h, *P*<0.05; ### = ionomycin 1 *μ*M 1 h *vs* ionomycin 1 *μ*M 24 h, *P*<0.001; #### = ionomycin 10 *μ*M 1 h *vs* ionomycin 10 *μ*M 24 h, *P*<0.0001). Statistical significance calculated by a 2-way ANOVA analyses with Tukey’s multiple comparison *post hoc* test. (**B ii**): Representative images of 24 h cultured untreated erythrocytes (top row), following treatment with PMA (10 *μ*M) (second row), ionomycin (10 *μ*M) (third row) and heat-shock treated (37°C) (bottom row). GFP channel (left), brightfield (centre) and merge (right). SYTOX Green (1 *μ*M; 20 minutes) was added after culture and images taken under 10x or 20x objective lens and throughout with the EVOS Auto FL2 microscope. FEETs formations (white arrows) and non-ETotic cell death (yellow arrows) indicated. Scale bars = 10 *μ*m. (**B iii**): Representative images of human neutrophils releasing NETs upon stimulation with PMA (10 nM) and ionomycin (5 *μ*M); 3 h culture in RPMI+5%FBS media. Scale bars = 10 *μ*m. (**C**): Time lapse images of erythrocytes cultured for 24 h with ionomycin (10 *μ*M) and treated with DNase-I (100 Units/mL). Cells stained with SYTOX Green (1 *μ*M; 20 min) and imaged with EVOS Auto FL2 microscope under 40x objective lens. Scale bars = 10 *μ*m.

To confirm the chromatin composition of FEETs, we incubated them with DNase-I and found that most of FEET-associated extracellular chromatin was digested within 90s (**Figure 1C; Figure S1, videos A and B**), thereby confirming the classical chromatin composition of FEETs. Next, we investigated the potential mechanism(s) involved in FEETs release, particularly whether PMA-induced FEETs were released upon PKC activation and in a NADPH oxidase dependent manner, as demonstrated in human NETs^21,25^. To investigate these pathways, erythrocytes were pre-treated with the pan-PKC inhibitor Ro 31-8820 and the NADPH oxidase inhibitor DPI, followed by PMA stimulation. DPI pre-treatment did not significantly prevent FEET release (**Figure S2 A**). Interestingly, after 48h, FEET release was substantially reduced when pre-treated with Ro 31-8820 (10*μ*M) ± PMA (**Figure 2A**). Therefore, we propose that PKC is involved in the PMA-induced FEETs pathway. To test the cellular pathways involved in ionomycin-induced FEETs, we pre-treated erythrocytes with the mitochondrial ROS (mROS) uncoupler DNP and the SK channels inhibitor apamin, both known to be integral for human NET production^22^. Apamin did not have any significant inhibitory effect (**Figure S2 B**), however, we noted that DNP pre-treatment significantly reduced FEETs formation, following 48h incubation with ionomycin (**Figure 2A**). Therefore, we conclude that mROS generation is involved in the ionomycin-induced FEETs pathway.

**Figure 2.**
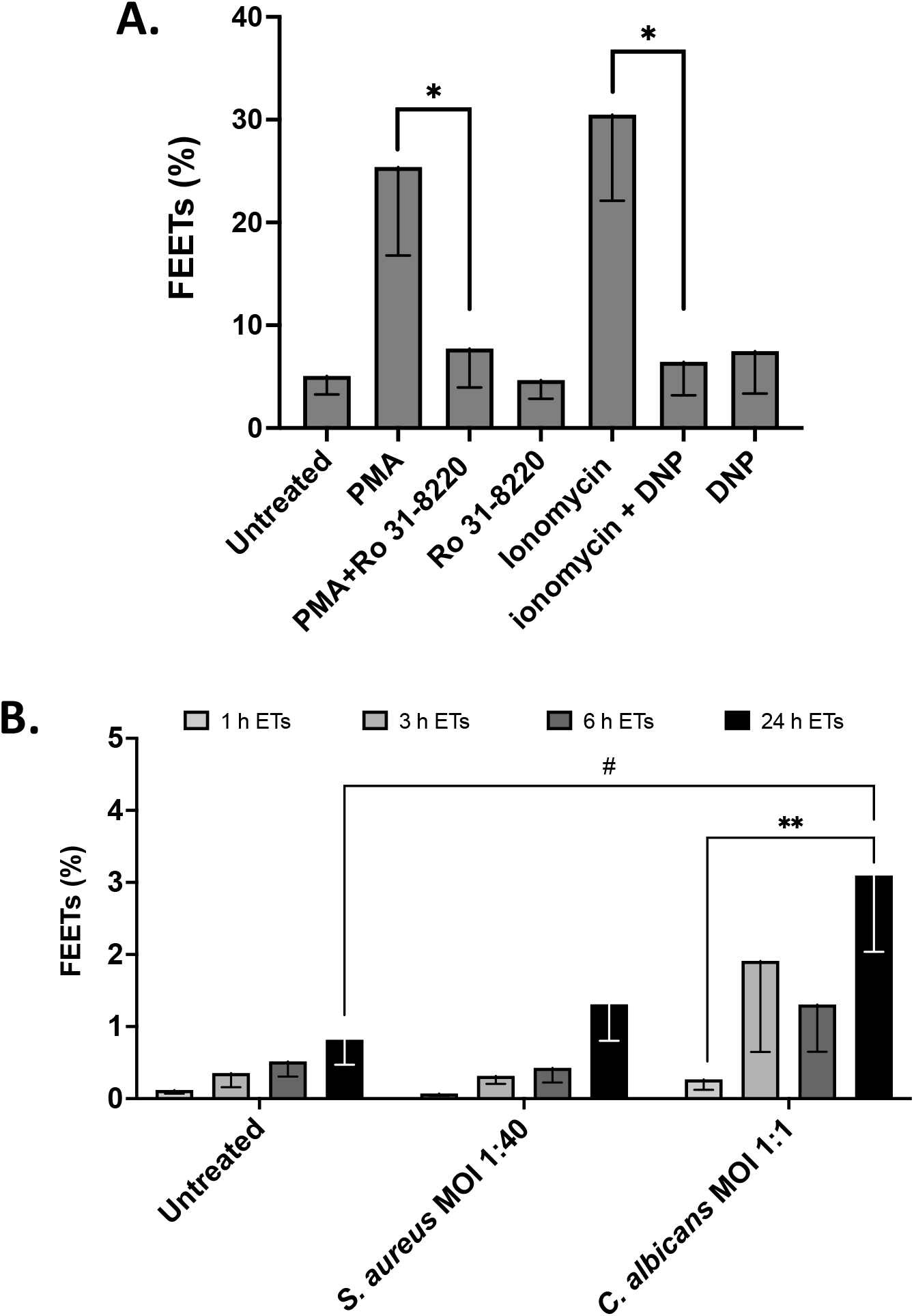
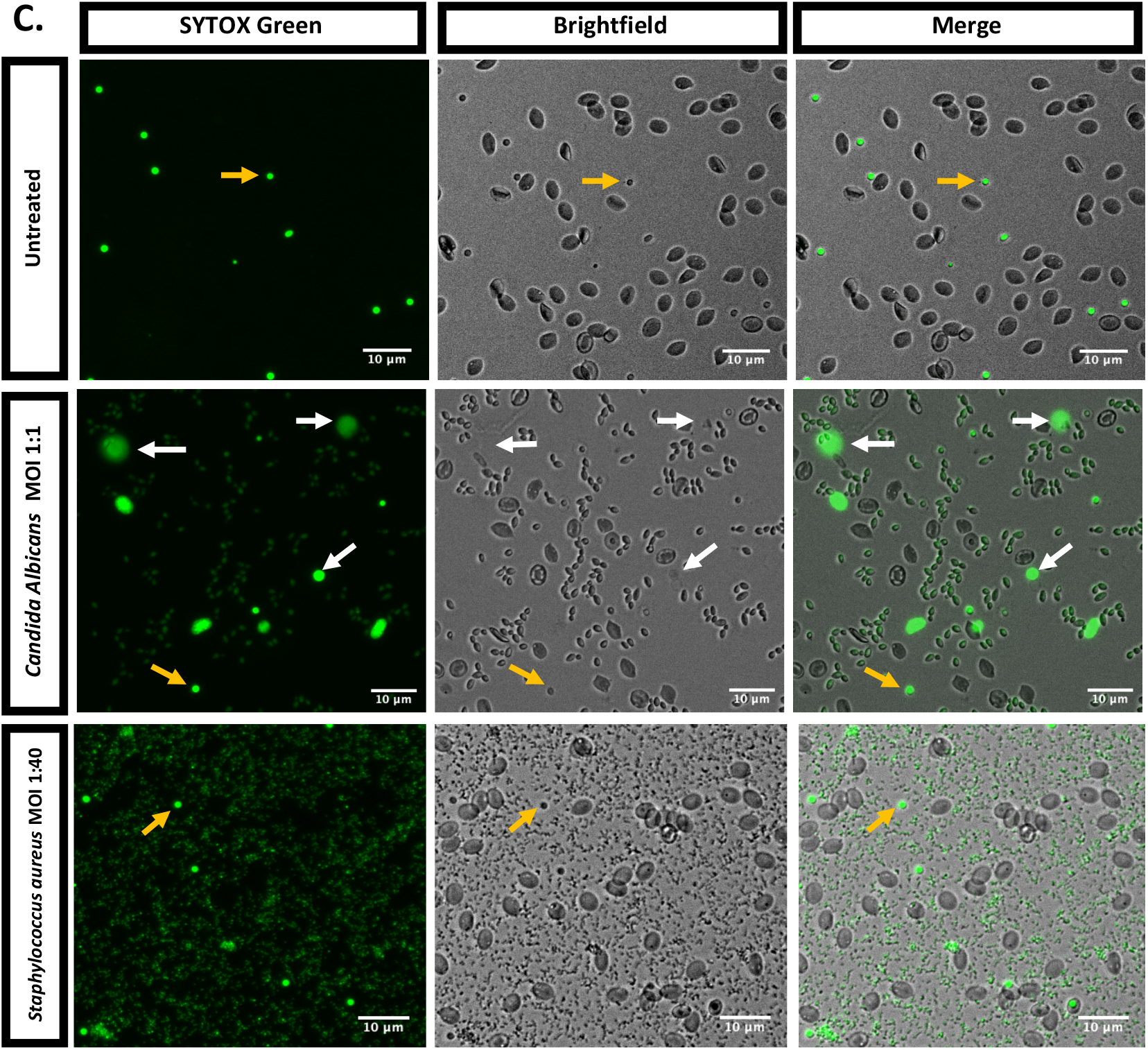
Zebrafish FEETs are released upon activation of PKC and generation of mitochondrial ROS and in response to physiological stimuli. (**A**): FEETs formations following treatment with PMA (10 *μ*M) or ionomycin (10 *μ*M) for 48 h ± prior treatment with Ro 31-8220 (10 *μ*M, 30 min) or DNP (187 *μ*M, 30 min). Values expressed as mean percentage ± s.e.m., n=6. Significant differences between PMA and PMA+Ro 31-8820 (*= PMA (10 *μ*M) *vs* PMA+Ro 31-8820 (10 *μ*M), *P*< 0.05) and between Ionomycin and Ionomycin+DNP (*= Ionomycin (10 *μ*M) *vs* Ionomyin+DNP (187 *μ*M), *P*< 0.05). Statistical analyses calculated with Paired *t-test*. (**B**): Quantification of FEETs formation in erythrocytes co-cultured with the heat-killed fungus *C. albicans* (MOI 1:1) and heat-killed Gram-positive bacterium *S. aureus* for 1, 3, 6 and 24 h. Values calculated as mean percentage ± s.e.m., n=6. Significant differences between untreated (control) and erythrocytes co-cultured with *C. albicans* (# = untreated 24 h *vs C. albicans* 24 h, *P*<0.05) and between *C. albicans* treated erythrocytes at different time points (** = *C. albicans* 1 h *vs C. albicans* 24 h, *P*<0.01). Statistical significance calculated by 2-way ANOVA analysis with Tukey’s multiple comparison *post hoc* test. (**C**): Representative images of erythrocytes untreated (top row), or 24 h co-cultured with *C. albicans* (middle row) and *S. aureus* (bottom row). After culture, cells were stained with SYTOX Green (1 *μ*M; 30 min). GFP channel (left), brightfield (centre) and merge images (right). White arrows indicate diffuse FEETs formations, yellow arrows indicate non-ETotic cell death. Images were taken with EVOS Auto FL2 microscope under 20x objective lens. Scale bars = 10 *μ*m.

Having demonstrated that zebrafish erythrocytes release FEETs upon chemical stimulation, we next investigated the possible antimicrobial functions of FEETs. We challenged erythrocytes with models of Gram-positive and fungal pathogens, specifically *S. aureus* and t *C. albicans*, both of which were heat-killed to prevent uncontrolled growth in culture media. FEETs were produced in response to *C. albicans* but not significantly in the presence of *S. aureus*. Notably, following 24h fungal challenge (MOI 1:1) there was a significant increase in the percentage of FEETs *vs* untreated erythrocytes (**Figure 2B**), albeit to a lesser extent compared with chemical simulation (**Figure 1B i,2A**). Furthermore, we observed a significant increase in the percentage of FEETs over time in erythrocytes co-cultured with *C. albicans* (MOI 1:1) at 24h vs identical culture conditions after 1h (**Figure 2B**). Representative fluorescent images of *C. albicans*-exposed erythrocytes show classical diffuse ETotic morphology (**Figure 2C, merge, white arrows**). No ET-like structures were observed after erythrocyte co-culture with *S. aureus* (in all MOIs tested including 1:40).Taken together, our results demonstrate for the first time, a previously unknown but potentially important role for erythrocyte involvement in immunity and host defence.

These results indicate evidence of possible fish erythrocyte antimicrobial function via formation of FEETs in response to *C. albicans*, thus highlighting a potential role in host defence and immune responses to infection. Our findings also provide a possible new explanation for why the nucleus is retained in fish erythrocytes in contrast to mammalian that undergo enucleation. Fish filter water through their gills to obtain dissolved oxygen, however, this site represents an entry channel for a multiplicity of pathogenic microorganisms causing infection and inflammation. The presence of >98% erythrocytes in circulating fish blood capable of exhibiting FEETs represents a rapid frontline of host defence against microbial invaders. Hitherto, FEETs have remained undescribed and our findings shed new light on erythrocyte-mediated immune defence in fish, as well as the conserved evolutionary nature of the ETotic response.

## Acknowledgments

The authors would like to thank all the staff of the aquatic facility in the Queen’s Medical Research Institute, University of Edinburgh for the help in fish maintenance and especially Dr Carl Tucker. In addition, the authors thank the staff of the Niall Bromage Freshwater Research Unit (University of Stirling) for providing technical assistance and care of the Atlantic salmon stocks. We thank EPSRC/MRC Centre for Doctoral Training in Optical Medical Imaging, OPTIMA (EP/L016559/1) for funding GR, the Medical Research Council UK (MR/K013386/1) for funding CTR and AGR. NAdH, APD and AGR were funded in part by a grant awarded to APD and AGR equally by BBSRC and NERC under the Sustainable Aquaculture Initiative (grant reference: BBM026132/1).

## Authorship contributions

Contribution: CTR, APD and AGR conceived the project; GR and NAdH performed experiments and analysed the data; GR wrote the manuscript; all the authors reviewed the final draft and revised it for critical content.

## Disclosure of Conflict of Interest

The authors declare no conflicts of interest.

## Notes

### Competing Interest Statement

The authors have declared no competing interest.

### Summary of Updates

Typos were found and corrected

