## supplementary materials for "Nucleated Fish Erythrocyte Extracellular Traps (FEETs) release is an evolutionary conserved immune defence process"

### Supplementary figure legends

#### Supplementary figure 1- DNase-I digestion of extracellular chromatin released by zebrafish FEETs.

Erythrocytes were cultured for 24 h (Video A) and 4 hours (Video B) in L-15 media with Ionomycin (10 $\mu$M; 28 °C) (Video A) or heat shocked at 37 °C (Video B). After culture, the DNA dye SYTOX Green was added (1 $\mu$M; 20 min). DNase-I (100 Units/mL) was added to digest extracellular DNA, time lapse was recorded with subsequent images taken every 30 s for 5 min with the EVOS Auto FL2 microscope. Scale bars 10 $\mu$m.

#### Supplementary figure 2- Reductive effects of NADPH-oxidase and SK channel inhibition on PMA- and ionomycin-induced zebrafish FEETs.

Quantification of FEETs released by erythrocytes pre-treated (30 min) with DPI (A) and Apamin (B) at different concentrations followed by 24 h stimulation with PMA (10 $\mu$M) (A) and ionomycin (10 $\mu$M) (B). Post culture, SYTOX green was added (1 $\mu$M; 30 min) and differential counts performed on images taken with the EVOS Auto FL2 microscope. Values are expressed as mean percentage ± s.e.m., n=6. No significant differences were found by comparing PMA *vs* PMA+DPI (at all concentrations tested) and Ionomycin vs Ionomycin+Apamin (at all concentrations tested).

#### Supplementary figure 3- Atlantic salmon erythrocytes release FEETs upon chemical stimulation.

(A): Representative image of erythrocytes isolated from Atlantic salmon and stained with the DNA dye DAPI (300 nM DAPI (Invitrogen) diluted in PBS) and the CD235a antibody (ab212432; Abcam). (B): Quantification of fluorescence intensity of erythrocytes stimulated with or without (RPMI control) ionomycin (0.1 $\mu$M) after 1 h of culture. Values are expressed as mean fluorescence intensity ± s.e.m., n=3. Statistical significant difference between RPMI control and ionomycin treated cells ($P < 0.05$). Statistical analyses calculated with Paired t-test. (C): Representative images of erythrocytes treated with ionomycin and stained with SYTOX green. GFP channel (left), brightfield (middle) and merge (right). Scale bars = 100 µm. Images were taken with an EVOS FL cell imaging system.

### Supplementary methods

#### Cytocentrifuge preparation

Whole zebrafish blood (100 $\mu$L, 1x$10^5$ cells) was cyto-centrifuged (300 rpm;3 min), fixed/permeabilised with methanol and stained with Diff-QuickTM (Gamidor) red and blue sequentially (Romanowsky stain variant). Each slide was mounted using DPX mounting media (Sigma) and imaged with a brightfield microscope.

#### Atlantic salmon erythrocytes collection and stimulation

Erythrocytes (red blood cells, RBCs) were isolated from Atlantic salmon blood collected aseptically from the caudal vein using a 2.5-mL syringe and 25G × 5/8 needle (Terumo, UK). Blood was diluted 5-fold in heparinized RPMI-1640 medium and loaded onto a 54% Percoll layer. RBCs were collected from the bottom pellet, washed and resuspended in chilled (4 ℃) RPMI-1640. Cells were seeded to 4×$10^5$ cells/ml into 96-well cell culture plates (Cell+; Sarstedt, Nümbrecht, Germany) containing RPMI-1640+2% FBS (Thermo Fisher Scientific) and supplemented with 1% penicillin-streptomycin (Sigma). Cells were allowed to settle for 30 min at 15 ℃ before exposure to 0.1 $\mu$M ionomycin (Sigma). Plates were incubated at 15 ℃ for 1 h before staining with SYTOX Green (5 $\mu$M, 20 min; Invitrogen). Fluorescence of each well was quantified with a microplate reader (Biotek Synergy HT, USA). Images were acquired with an EVOS FL cell imaging system (Thermo Fisher Scientific) equipped with bright field, 357/44 nm and 470/22 nm excitation LED lights. Images were captured with a ×20 and ×40 objectives, and processed using the NIS-Elements 3.2 software.

#### Atlantic salmon immunofluorescence staining

To assess the purity of the red blood cell population, the specific erythrocyte membrane marker CD235a was probed with a mouse monoclonal antibody raised to human glycophorin A (Abcam, ref. 212432). RBCs incubated in 96-well cell culture plates were fixed with 4% paraformaldehyde in PBS (20 min, RT). After fixing, cells were washed with PBS and blocked with 3% bovine serum albumin (BSA) in PBS (60 min). Then, the primary antibodies were added at 1:200 dilution and incubated overnight at 4ºC. After three washes with PBS, conjugated Alexa Fluor 488 (Invitrogen) diluted 1:300 in PBS supplemented with 3% BSA was added (90 min). Cells were washed thrice with PBS to remove excess antibodies and then incubated with 300 nM DAPI to counterstain nuclei. Images were acquired as described previously.

### Supplementary figures

Supplementary figure 1 – *DNase-I digestion of extracellular chromatin released by zebrafish FEETs.*

(Videos attached separately)

Supplementary figure 2 – *Reductive effects of NADPH-oxidase and SK channel inhibition on PMA- and ionomycin-induced zebrafish FEETs.*

**A.**

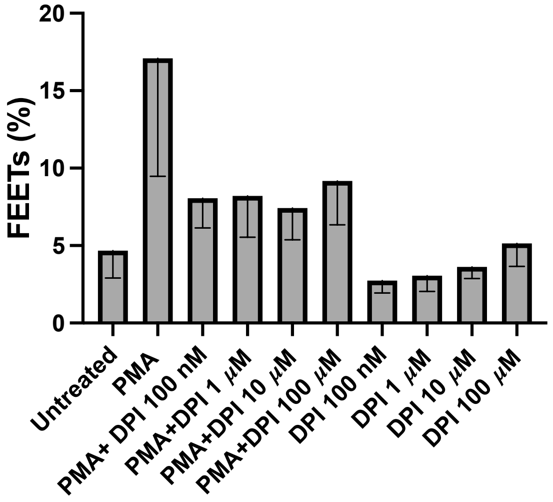


**B.**

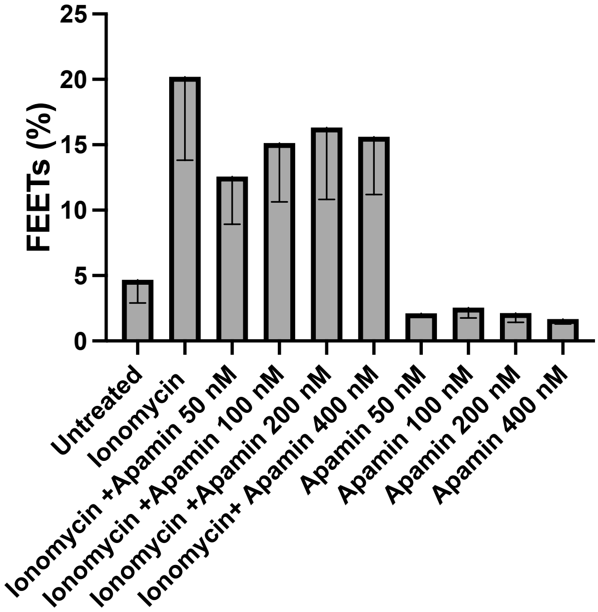

Supplementary figure 3 – *Atlantic salmon erythrocytes release FEETs upon chemical stimulation.*

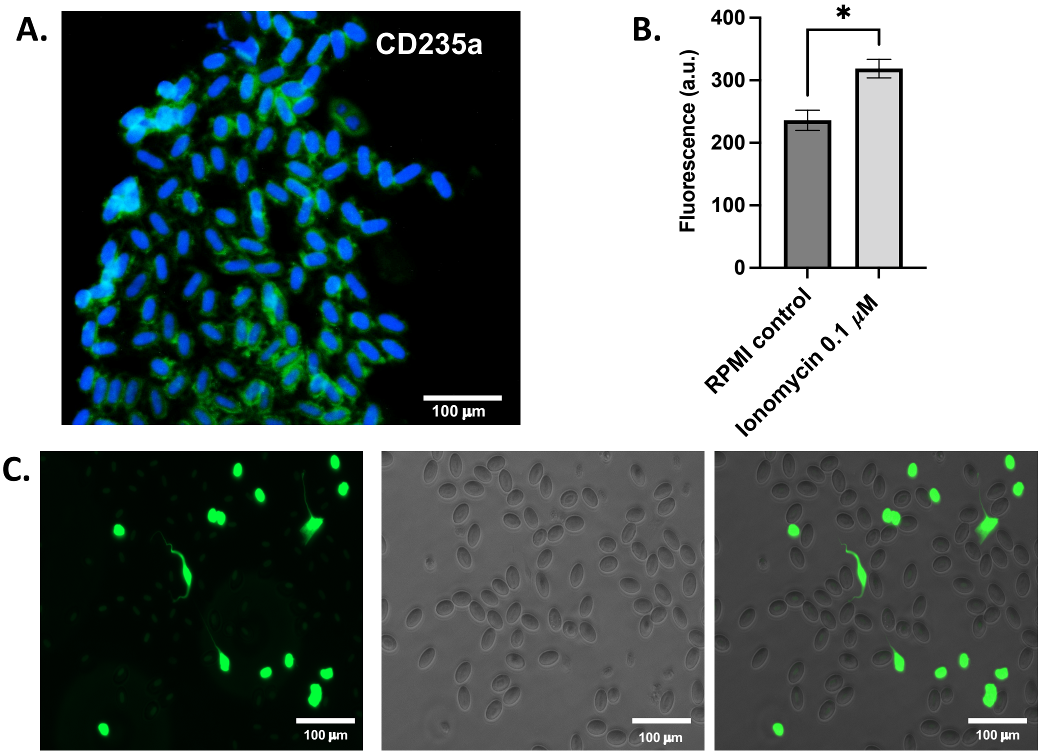
